# Promoting social connectedness through multi-person neurofeedback

**DOI:** 10.1101/2025.05.22.655644

**Authors:** Xiaojun Cheng, Rongbin Zhang, Phoebe Chen, Ziyuan Song, Feng Cheng, Suzanne Dikker, Yafeng Pan

**Author notes:** Shared senior authorship, correspondence should be addressed to Yafeng Pan, Department of Psychology and Behavioral Sciences, Zhejiang University, Hangzhou, China.; Suzanne Dikker, Department of Experimental Psychology, Ghent University, Ghent, Belgium.

## Abstract

Humans are inherently driven to build meaningful relationships, but attempts to socially connect with others are not always successful. This study investigates whether social connectedness can be improved by intentionally regulating inter-brain coupling, a neural correlate of successful social interactions. Using a multi-person neurofeedback system (i.e., a multi-brain computer interface), we showed dyads real-time visualizations of the extent to which their brainwaves (EEG signals) were “in sync”. Our results showed that, compared to a sham control group, dyads who received multi-brain neurofeedback exhibited an increase in inter-brain coupling, and, critically, that this increase was associated with a higher sense of social connectedness. A chain mediation analysis suggested that this experience of enhanced social connectedness may have been driven by a sense of joint control and shared intentionality. Together, our findings showcase the potential for regulating inter-brain coupling to optimize human social relationships and behaviors.

## 1. Introduction

Social connectedness is essential for our psychological and physical well-being: it provides us with a sense of identity, purpose, and belonging ^1,2^, and the lack of rewarding social interactions has been linked to adverse health effects ^3,4^. Individuals who perceive a strong sense of social connectedness in general often experience deep emotional bonds with others, readily empathize with them, view them as approachable and friendly, and actively engage in social groups and activities ^5,6^. In recent years, studies have linked perceived social connectedness with the extent to which brain activity becomes coupled during social interactions, such as interpersonal coordination and sharing emotional experiences ^7,8^. Inter-brain coupling has been found to correlate with interaction quality across various activities, including interpersonal communication ^9,10^, interactive learning ^11,12^, and economic decision-making ^13,14^. Notably, inter-brain coupling during interactions has been linked to subsequent bonding and social behaviors such as helping and cooperating ^13,15^. These findings raise the question if inter-brain coupling can be externally induced, and if so: if it can be leveraged to create interventions to improve social connectedness.

Indeed, recent studies have observed that induced inter-brain coupling is predictive of dyads’ empathy ^16^, interpersonal coordination ^17^, and learning performance ^11^. These findings suggest that changes in inter-brain coupling can affect subsequent cognitive and affective states. One potential explanation is that inter-brain coupling might directly impact the internal shared mental processes between individuals during an interaction ^16,18,19^. Shared mental processes refer to the alignment of cognitive and emotional processes between individuals, encompassing various psychological activities associated with “theory of mind”, such as shared intentionality and perceived similarity ^20–22^. Shared intentionality involves a mutual recognition of goals and actions within a group ^23^, while perceived similarity highlights the commonality in how individuals perceive their companions as similar to themselves, reflecting a shared sense of how the world is experienced and interpreted ^24,25^. These shared mental processes are closely related to social connectedness ^26,27^. In this study, we investigated whether shared intentionality and perceived similarity would mediate the effect of endogenous regulation of inter-brain coupling on social connectedness.

In recent years, neurofeedback has emerged as a promising intervention for regulating brain activity in individuals ^28,29^. During neurofeedback interventions, participants interact with real-time visual, auditory, or haptic representations of their brain activity using a brain-computer interface (BCI). Typically, the BCI targets specific neural indices (e.g., alpha power) with the goal to help improve a specific cognitive function (e.g., attention) ^30–32^. Neurofeedback is commonly used in combination with electroencephalography (EEG) due to its high temporal resolution ^33,34^. For example, EEG-based neurofeedback has been used with the aim to help improve cognitive performance ^30,35^, sleep quality, and emotional control ^36^. Neurofeedback can also be extended to a group of individuals (i.e., multi-person neurofeedback, or cross-brain neurofeedback), providing feedback targeting indicators of *multiple* brains to enable dyads or group members to regulate inter-brain coupling ^37–40^. In most multi-brain biofeedback studies to date, participants were explicitly instructed to try out strategies to maximize their inter-brain coupling (“what does it mean to be on the same brain wavelength?”) and participants received feedback about their inter-brain coupling using a BCI that rewarded inter-brain coupling e.g., through light patterns and avatar sizes/distances. These studies suggest that multi-brain neurofeedback training may indeed lead to an increase in inter-brain coupling, implying the potential for active induction of inter-brain coupling through external training ^38,40^. While these findings are promising, they also leave some questions unaddressed. Most notably, it is unclear under which conditions multi-person neurofeedback can help improve social connectedness. For example, the observed increases in inter-brain coupling may arise from the awareness that the BCI reflects their inter-brain coupling. Alternatively, participants may actually be able to extract meaningful information from the BCI directly, effectively manipulating synchrony with their collective behavior. And if participants experience a sense of control, then what might be the associated internal mental processes that allow them to exert such control?

Here, we explicitly investigated the interplay between multi-brain neurofeedback and social connectedness. Fifty-seven dyads underwent a 12-minute multi-person neurofeedback, during which they imagined synchronizing their brain activity with each other. The BCI consisted of two line-drawn brains approaching each other or separating from each other based on the level of synchronization ^37^. Inter-brain coupling was quantified in real time using online coherence of EEG signals (see Methods). Critically, half of the dyads were presented with a BCI based on real neural signals and half were shown a BCI based on randomly generated signals (real-NFB vs. sham-NFB). Both groups received the same instruction that the BCI would reflect their inter-brain coupling; this allowed us to test whether the realness of the data would bias their sense of joint control. Before and after the imagery task, participants completed subjective measurements evaluating their experience of social connectedness and shared mental processes. We hypothesized that:

*H1: Multi-person neurofeedback impacts inter-brain coupling (more inter-brain coupling during the task, but only in the real-neurofeedback group);*

*H2: Multi-person neurofeedback can promote social connectedness (dyads in the real-NFB group report a higher increase of social connectedness than the sham-NFB group)*;

*H3: The experience of increased social connectedness is linked to shared mental processes (i.e., shared intentionality and perceived similarity)*.

## 2. Methods

### 2.1. Participants

A total of 114 college students (36 males and 78 females, aged 20.08 ± 2.19 years, mean ± standard deviation) were recruited for the study. They were randomly assigned to 57 dyads, with 28 dyads in the real-NFB group and 29 dyads in the sham-NFB group. To mitigate the effects of gender and familiarity ^41^, each dyad consisted of two participants of the same gender who were previously unacquainted with each other. All participants were right-handed, had normal or corrected-to-normal vision, and had no history of neurological or psychiatric disorders. Prior to the experiment, each participant provided informed consent and they were compensated with 60-70 yuan upon completion. This study was approved by the Ethical Institute Review Board of Shenzhen University.

### 2.2. Experimental tasks and procedures

Two participants were seated next to each other (**Figure 1A**), in a quiet laboratory in front of a screen (DELL, 24-inch, 1080p, 60Hz). Each dyad was equipped with a portable wireless EEG headset (EMOTIV EPOC X) (**Figure 1B**), for real-time computation of inter-brain coupling and visual feedback (**Figure 1C**).

**Figure 1.**
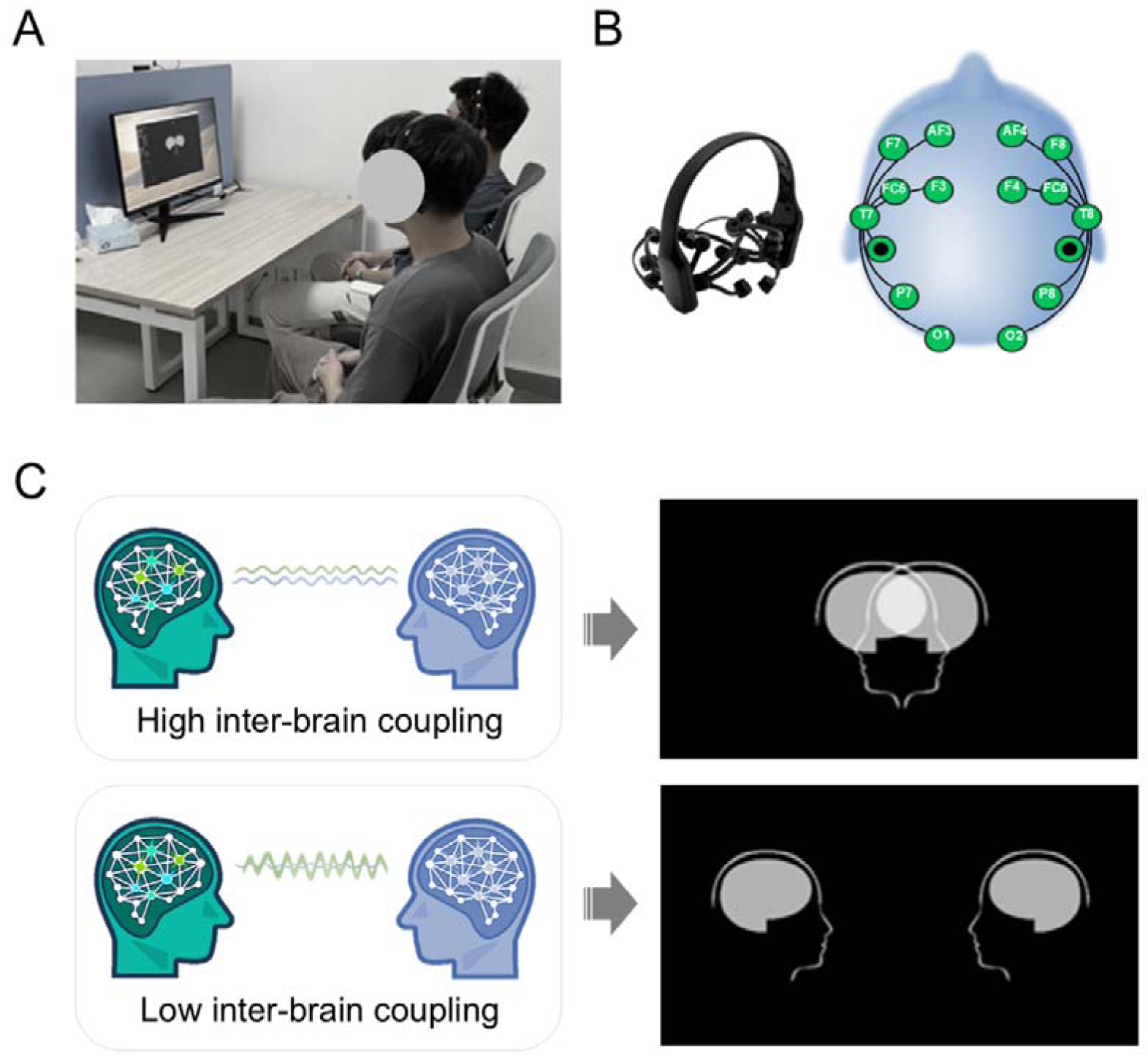
Multi-person neurofeedback setup. (A) Experimental setup. (B) Portable EEG device and electrodes configuration. (C) The overlap of the two avatar icons represents a higher inter-brain coupling between the participants, while their separation indicates a lower inter-brain coupling.

During the experiment, dyads underwent four successive sessions: a pre-training test session, a 4-minute resting-state session, a 12-minute multi-person neurofeedback training session, and a post-training test session (**Figure 2**). The details of each session are outlined below.

**Figure 2.**
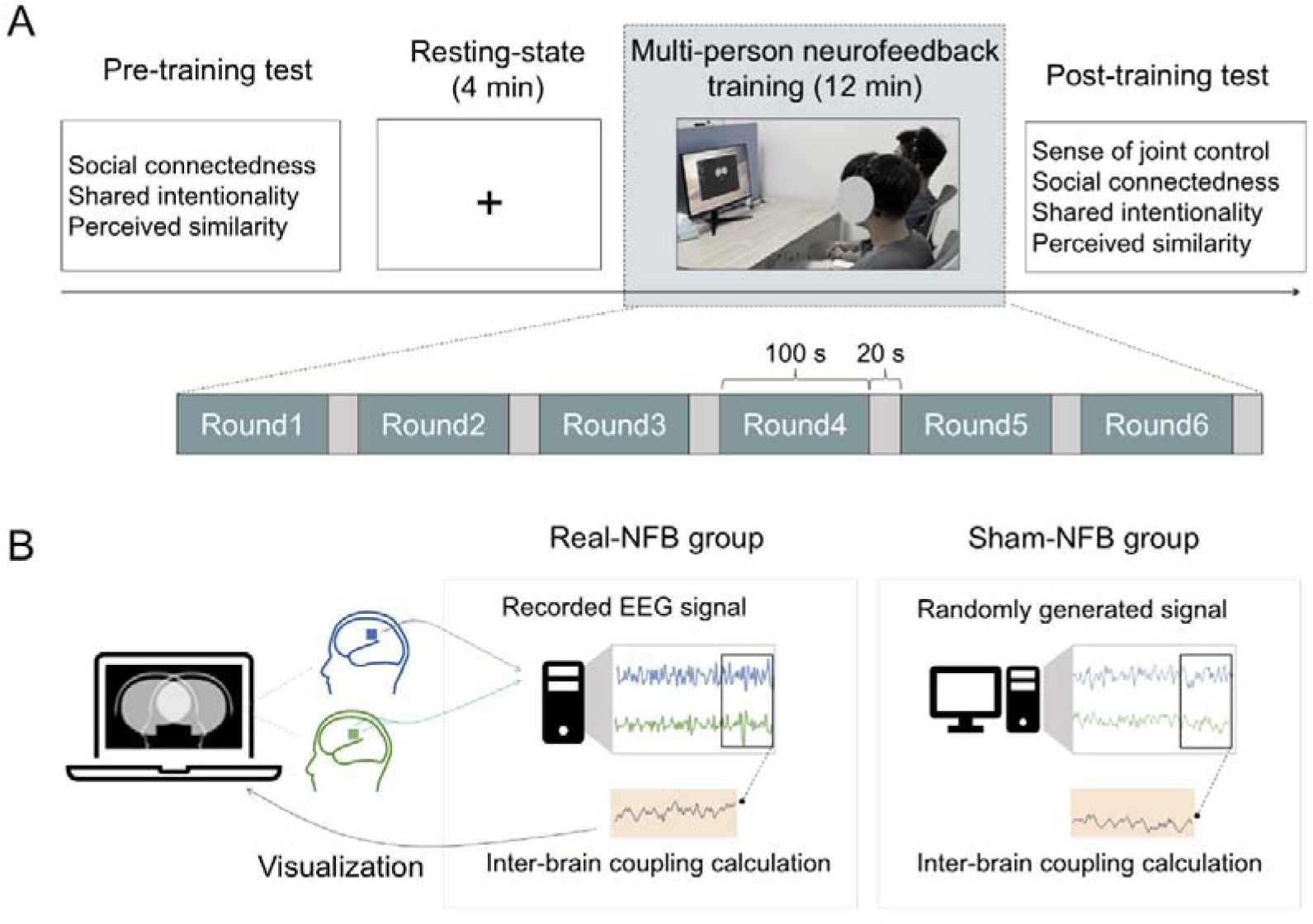
Experimental design. (A) Experimental procedure. (B) Participants were randomly assigned to the real-neurofeedback (NFB) group or a sham control group. In the real-NFB group, feedback was delivered according to the synchronization computed from the actual EEG signals of the participants. Conversely, in the sham-NFB group, feedback was based on the synchronization calculated from data generated by the program. (C) Participants completed six rounds of multi-person neurofeedback training, with each round lasting 100 seconds and a 20-second break between rounds.

#### 2.2.1. Session I: Pre-training test session

Participants were asked to individually complete questionnaires measuring social connectedness and shared mental processes. During this session, no verbal or nonverbal communication was allowed.

##### Social connectedness

Based on previous studies ^42,43^, four items from the emotional bonding questionnaire were extracted to evaluate participants’ feelings of social connectedness, including psychological closeness (i.e., "How close do you feel to the partner?"), friendliness (“To what extent do you feel you could be friends with the partner?”), likability (“How much do you like the partner?”), and trust towards their partner to their partner (“How much do you trust the partner?”) ^44,45^. All items were rated on a 9-point Likert scale, ranging from 1 (not at all) to 9 (strongly). The results from the four items were averaged.

##### Shared mental processes

To evaluate participants’ shared mental processes, we used two sets of questionnaires to examine *shared intentionality* and *perceived similarity* during the task. Regarding *shared intentionality,* five items were drawn from the rapport questionnaire ^26,46^. Participants responded to statements such as: “During interaction with my partner, there was a shared flow of thoughts and feelings”. For *perceived similarity,* five items were used ^47^, with participants rating their degree of similarity to other participants. All questionnaire items were rated on a 9-point Likert scale, ranging from 1 (not at all) to 9 (strongly). The results from the items for shared intentionality and perceived similarity were averaged.

#### 2.2.2. Session II: Resting-state session

Upon completing the pre-experiment tests, participants engaged in a 4-minute resting-state session. During this session, participants were asked to remain still, relax their minds, and refrain from focusing on specific issues. The first two minutes involved closing their eyes, followed by two minutes with their eyes open ^40^.

#### 2.2.3. Session III: Multi-person neurofeedback training session

Then, participants underwent a multi-person neurofeedback training task. In this task, they viewed a BCI display featuring two avatar icons, with instructions that each represented their own brain. They were told that the proximity of the two icons on the screen varied based on the synchronization level of their brain activity; closer proximity indicated greater synchronization. In both groups, participants were asked to concentrate on pushing the icons closer together with their minds, aiming to synchronize their brain activity with their partner’s. Example instructions (provided in Chinese) include “*During the task, you will see two avatar icons on the screen representing your brains. Your goal is to push these avatars together with your minds so they overlap in the center of the screen. The icons will change in real time based on the synchronization between your brainwaves*”. To achieve this, they needed to adjust their brainwaves to align with their partner’s as closely as possible. This was an intentionally abstract instruction that was supposed to help the participants use different strategies. Consistent with recent practice ^40^, participants were not provided with any instructions regarding the different experimental conditions (real-NFB vs. sham-NFB), ensuring that they remained blind to the testing conditions. Only nonverbal interactions were allowed during this session, as verbal communication is more likely to introduce additional cognitive and emotional layers (e.g., language processing and emotional tone), which could complicate the interpretation of the neurofeedback. Each dyad completed six rounds of multi-person neurofeedback, each lasting 100 seconds, with a 20-second break between rounds (**Figure 2A**). In the real-NFB group, dyads were presented with a BCI based on real-time neural signals; in the sham-NFB group, dyads were shown a BCI based on the synchronization calculated from random data (i.e., random samples drawn from a uniform distribution between 0 and 1) generated by the Hybrid Harmony program ^37^.

During the task, the raw EEG signals of dyads were recorded using two 14-electrode portable wireless EEG headsets (EMOTIV EPOC X), with a sampling rate of 128Hz. The LabRecorder program was used to save data through Lab Streaming Layer (LSL; https://github.com/sccn/labstreaminglayer). EEG data streams from the two participants’ headsets were synchronized using a 3-second window, based on the time that samples were received by the analysis computer, and then filtered into typical frequency bands: delta (1-3 Hz), theta (4-8 Hz), alpha (9-12 Hz), and beta (13-30 Hz). Real-time online inter-brain coupling computations were provided to dyads via the *Hybrid Harmony* program ^37^, running on Python 3.6. Online inter-brain coupling between individuals was computed using Coherence between EEG signals from all one-to-one electrode pairs (e.g., Fp1 is only correlated with Fp1 and so on) every 3-second epoch ^48^, and then averaged across all homologous pairs of electrodes between headsets and across all epochs. The inter-brain coupling values for each frequency were then fed into a visualization algorithm ^37^: higher inter-brain coupling values brought the avatar icons closer together until they completely overlapped; conversely, lower inter-brain coupling values resulted in the icons moving farther away (**Figure 1C**). The overview and full protocol of implementing multi-person neurofeedback has been released publicly (https://github.com/RhythmsOfRelating/HybridHarmony).

#### 2.2.4. Session IV: Post-training test session

Following the multi-person neurofeedback training, participants were asked to complete post-training surveys, which included questionnaires measuring social connectedness and shared mental processes (same as pre-training tests). They also reported their sense of joint control during the task. Specifically, they responded to two questions regarding their perceived control over the movement of their avatar icon: “To what extent do you feel you can control the movement of your avatar icon?” and “To what extent do you feel you can control the movement of the avatar icon together?” Responses were rated on a 9-point Likert-type scale, ranging from 1 (did not feel in control at all) to 9 (felt totally in control). A mean score of the sense of control was computed based on those two questions.

### 2.4. Data analysis

#### 2.4.1. Behavioral data

For all behavioral data and subjective reports, the scores of participants in dyads were averaged to represent dyad-level scores. Repeated-measures ANOVAs were conducted on social connectedness and shared mental processes, with Group (real-NFB vs. sham-NFB) as the between-dyad variable and Time (pre-training vs. post-training) as the within-dyad variable. For sense of joint control, independent-samples *t*-tests were performed on post-training data, with Group as the between-dyad variable. Additionally, Pearson correlation analysis was conducted to investigate the relationship among social connectedness, shared mental processes, and sense of joint control. Specifically, we focused on the increase of social connectedness (ΔSocial connectedness) and shared mental processes (ΔShared intentionality, ΔPerceived similarity) from the pre-training to post-training when performing correlation analysis.

#### 2.4.2. EEG data

Our EEG data underwent both online and offline analyses. We used the Coherence metric to compute online inter-brain coupling ^48^. Coherence is computationally simple and fast to compute, making it suitable for real-time applications where quick feedback is essential. Real-time neurofeedback systems require immediate calculations to provide instantaneous feedback to participants. For real-time visualization purposes, online inter-brain coupling computation underwent a [0 0.3] min-max normalization ^37^. In this online analysis, coherence in the sham group was calculated using randomly generated signals, which were used to deliver visual feedback to the dyads, rather than the actual signals.

To parse true signals from both groups, we computed inter-brain coupling using circular correlation coefficient (CCorr) analysis over the data post-hoc (offline analysis). CCorr is commonly used to compute inter-brain coupling in hyperscanning studies to capture phase relationships, in part because it is less sensitive to spurious correlations than some other metrics ^49–52^. CCorr is less suitable for real-time analyses because of computational constraints that can induce temporal lags in the neurofeedback (aka. reducing the “real-time” nature of the experience). Offline analysis also provides the opportunity to preprocess the data more thoroughly, including the removal of artifacts and noise that might affect the calculation of inter-brain coupling. This allows for the application of circular correlation to cleaner data, yielding more accurate and reliable results. To preprocess EEG data for the offline analysis, we used EEGLAB on MATLAB R2020a (MathWorks Inc., USA). Bad channels were excluded and an average re-reference was applied to the raw EEG data^40^. A high-pass filtering at 0.5 Hz was performed to eliminate slow fluctuations ^38^. We performed Independent Component Analysis (ICA) and then used the ADJUST toolbox implemented in EEGLAB to semi-automatically identify artifacts related to heart rate and eye blink ^53,54^. The effectiveness of ICA on EMOTIV devices has been previously validated ^55^.

Phase values were extracted from the preprocessed data using the Morlet wavelet transform on signals ^13,56^ from 1 to 30 Hz, with a step size of 1 Hz. We calculated CCorr by using the Circular Statistics Toolbox to evaluate dyadic inter-brain couplings ^50,52,57^. CCorr measures the covariance between the phase time series of two neural signals. Specifically, it quantifies how consistently the phase of one signal changes in relation to the phase of another signal over time. Its function is defined as:

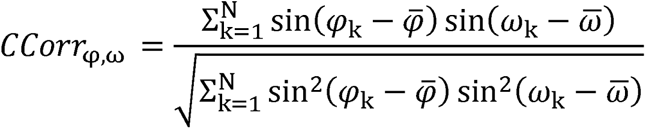

where _φ_ and _ω_ represent the phase values of the two signals, while 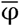 and 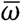 denote the mean phases of these signals. Following previous recommendations ^57^, the absolute CCorr was used as the indicator of inter-brain coupling. This coefficient represents the absolute value of the Fisher-*z* transformation of the circular correlation between two time series signals, i.e., inter-brain coupling = |zCCorr|.

Throughout the six rounds of the multi-person neurofeedback task, we discarded the last 10 seconds of each round to mitigate edge artifacts. The remaining data were used to compute task-related inter-brain coupling. First, we calculated the CCorr of every 3-second interval across all possible electrode combinations (14 * 14) for each dyad and frequency ^57^. This was done for both the neurofeedback session (Session III) and the resting-state session (Session II). Then, inter-brain coupling of each group was quantified by averaging CCorr values across all possible electrode combinations and time windows for each frequency ^38^. Task-related inter-brain coupling was calculated by subtracting inter-brain coupling at the resting-state session from inter-brain coupling at the multi-person neurofeedback session. In the remainder of the paper, we refer to task-related inter-brain coupling as inter-brain coupling. These steps ultimately yielded 30 inter-brain coupling values ranging from 1 to 30 Hz for each dyad.

Following inter-brain coupling computation, a frequency-to-frequency independent sample *t*-test was performed to compare the effects of different multi-person neurofeedback training groups (real-NFB vs. sham-NFB), with multiple comparisons correction applied using False Discovery Rate (FDR) at a threshold of *p* < 0.05. Additionally, Pearson correlation analysis was conducted to investigate the relationship between inter-brain coupling and social connectedness, as well as shared mental processes. Specifically, we focused on the difference in subjective measurements (scores of social connectedness and shared mental processes) between the post-training tests and the pre-training tests (ΔSocial connectedness, ΔShared intentionality, and ΔPerceived similarity).

## 3. Results

### 3.1. Validation of multi-person neurofeedback

As a first-pass check, we examined whether multi-person neurofeedback training successfully induced inter-brain coupling. As expected, the real-NFB group demonstrated significantly higher inter-brain coupling than the sham-NFB group, *t* (25.65) = 4.979, *p* < 0.001, Cohen’s *d* = 1.395. This indicates that the two groups experienced different proximities of the two icons on the screen (real-NFB: relatively closer, sham-NFB: relatively farther).

For the offline analysis, frequency-wise independent sample *t*-tests revealed that inter-brain coupling in the real-NFB group was significantly higher than that in the sham-NFB group at 21-23 Hz (uncorrected *p*s < 0.05; **Figure 3A**). Inter-brain coupling at 22 Hz survived after FDR correction (corrected *p* = 0.045). We then computed the averaged inter-brain coupling for 21-23 Hz, and found that inter-brain coupling at this frequency was higher in the real-NFB group than that in the sham-NFB group, *t* (55) = 2.931, *p* = 0.005, Cohen’s *d* = 0.857. In the remainder of the manuscript, we focused on the offline inter-brain coupling values.

**Figure 3.**
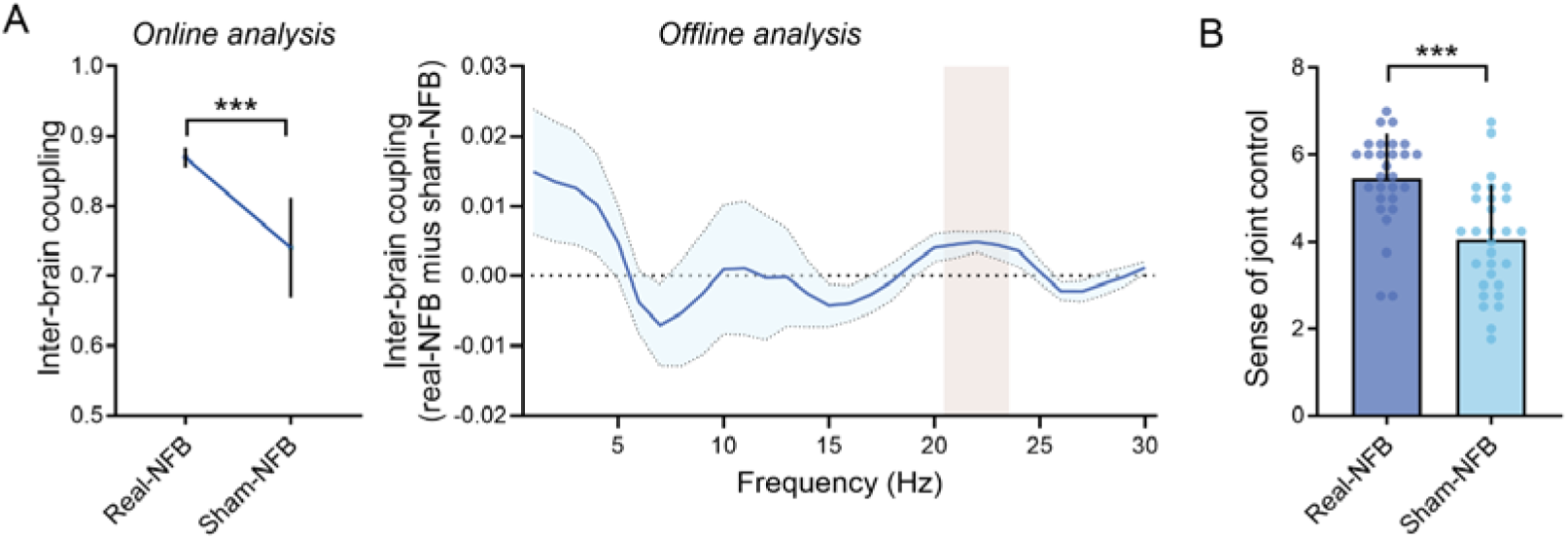
Validation of multi-person neurofeedback (NFB). The real-NFB group elicited enhanced task-related inter-brain coupling in the online analysis (A, left panel), and in the offline analysis at 21-23 Hz, highlighted in a vertical bar (A, right panel), and reported a significantly higher sense of joint control (B), compared to the sham-NFB group. * *p* < 0.05, *** *p* < 0.001. Shadows around the means and error bars denote standard deviations.

Complementary to the above validation, we also assessed the training effects on the sense of joint control. Participants in the real-NFB group reported a significantly higher sense of joint control compared to the sham-NFB group (**Figure 3B**), *t* (55) = 4.542, *p* < 0.001, Cohen’s *d* = 1.203. These results indicate that multi-person neurofeedback in the present study successfully induced stronger inter-brain coupling, and participants in the two groups perceived different levels of the sense of joint control. These findings support our confirmatory hypothesis H1.

### 3.2. Effects of multi-person neurofeedback on social connectedness and shared mental processes

Having confirmed that multi-person neurofeedback successfully regulated inter-brain coupling, we next tested our hypothesis H2, evaluating the effects of neurofeedback on social connectedness. The results showed that there was a significant main effect of Time (pre-training vs. post-training), *F* (1, 55) = 18.992, *p* < 0.001, η^2^_partial_ = 0.257, with enhanced social connectedness after neurofeedback. No main effect of Group (real-NFB vs sham-NFB) was found, *F* (1, 55) = 2.646, *p* = 0.110, η^2^_partial_ = 0.046. More importantly, we found a significant interaction effect, *F* (1, 55) = 5.695, *p* = 0.020, η^2^_partial_ = 0.094, indicating that in the real-NFB group, the post-training session yielded a higher level of social connectedness compared with the pre-training session, *F* (1, 55) = 22.351, *p* < 0.001, η^2^_partial_ = 0.289, while such effect was absent in the sham-NFB group, *F* (1, 55) = 1.978, *p* = 0.165, η^2^_partial_ = 0.035 (**Figure 4A**). These findings provide confirmation that multi-person neurofeedback can indeed predict an enhancement in social connectedness, thereby supporting our hypothesis H2.

**Figure 4.**
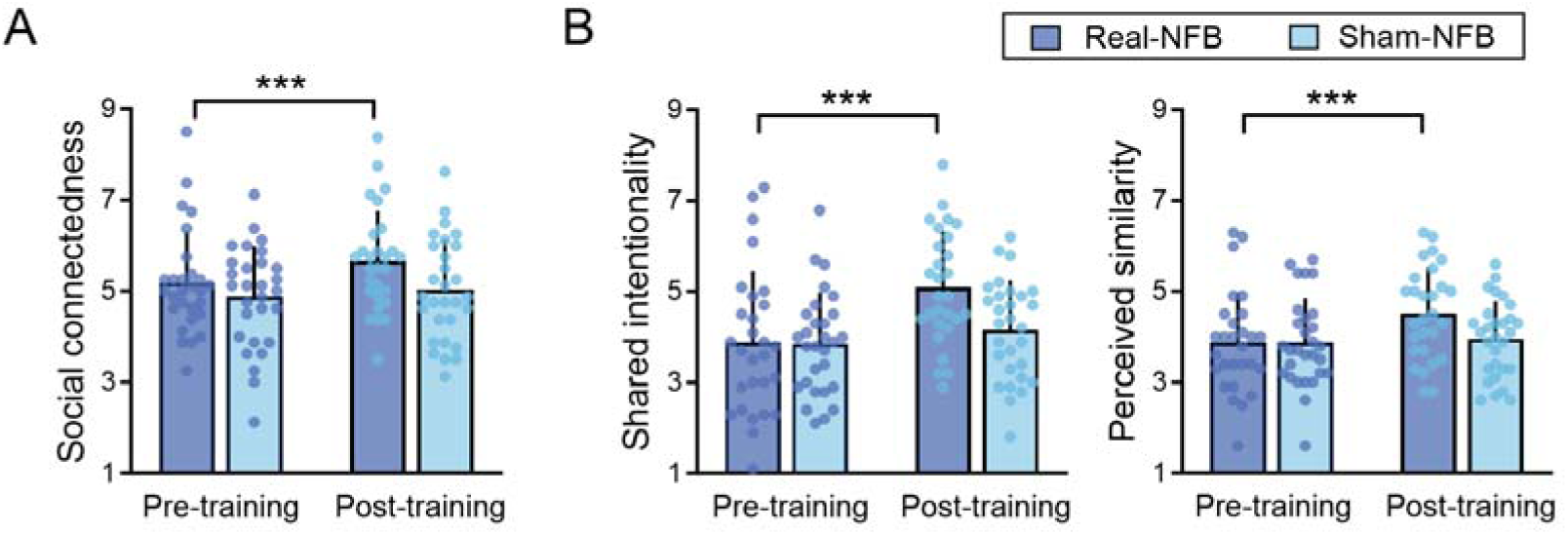
Effects of multi-person neurofeedback (NFB) on social connectedness and shared mental processes. (A) The real-NFB training promoted social connectedness. (B) The real-NFB training promoted shared mental processes, including shared intentionality and perceived similarity. *** *p* < 0.001. Error bars denote standard deviations.

To test our hypothesis H3, we first assessed whether multi-person neurofeedback could likewise promote shared mental processes (including shared intentionality and perceived similarity). For shared intentionality, the results revealed a significant main effect of Time, *F* (1, 55) = 29.253, *p* < 0.001, η^2^_partial_ = 0.347, indicating a higher level of shared intentionality after neurofeedback training. No significant main effect of Group was detected, *F* (1, 55) = 2.438, *p* = 0.124, η^2^_partial_ = 0.042. Notably, a significant interaction effect was observed, *F* (1, 55) = 10.598, *p* = 0.002, η^2^_partial_ = 0.162. Specifically, in the real-NFB group, shared intentionality significantly increased at post-training compared to pre-training, *F* (1, 55) = 36.886, *p* < 0.001, η^2^_partial_ = 0.401, while such observation was absent in the sham-NFB group, *F* (1, 55) = 2.360, *p* = 0.130, η^2^_partial_ = 0.041 (**Figure 4B, left)**. Regarding perceived similarity, the main effect of Time was significant, *F* (1, 55) = 9.600, *p* = 0.003, η*^2^_partial_* = 0.149, showing that perceived similarity was enhanced after neurofeedback training. The main effect of Group was not significant, *F* (1, 55) = 1.487, *p* = 0.228, η*^2^_partial_* = 0.026. Moreover, there was a significant interaction effect on perceived similarity, *F* (1, 55) = 6.006, *p* = 0.017, η^2^_partial_ = 0.098. In the real-NFB group, perceived similarity significantly increased at post-training compared to pre-training, *F* (1, 55) = 15.130, *p* < 0.001, η*^2^_partial_* = 0.216, while no such effect was observed in the sham-NFB group, *F* (1, 55) = 0.213, *p* = 0.646, η^2^_partial_ = 0.004 (**Figure 4B, right)**. These findings suggest that multi-person neurofeedback also improved shared mental processes.

### 3.3. Multi-person neurofeedback-regulated inter-brain coupling affects social connectedness through the sense of joint control and shared mental processes

To explore the relationships among the multi-person neurofeedback-regulated inter-brain coupling, sense of joint control, shared mental processes, and social connectedness, we carried out a series of Pearson correlation analysis. We observed that inter-brain coupling at 21-23 Hz was associated with a sense of joint control, *r* = 0.292, *p* = 0.028 (**Figure 5A**). Further, a sense of joint control was correlated with Δ Shared intentionality, *r* = 0.531, *p* = < 0.001, with Δ Perceived similarity, *r* = 0.344, *p* = 0.009, and with Δ Social connectedness, *r* = 0.486, *p* < 0.001 (**Figure 5B**). More importantly, both Δ Shared intentionality (*r* = 0.557, *p* < 0.001) and Δ Perceived similarity (*r* = 0.457, *p* < 0.001) were correlated with Δ Social connectedness (**Figure 5C**).

**Figure 5.**
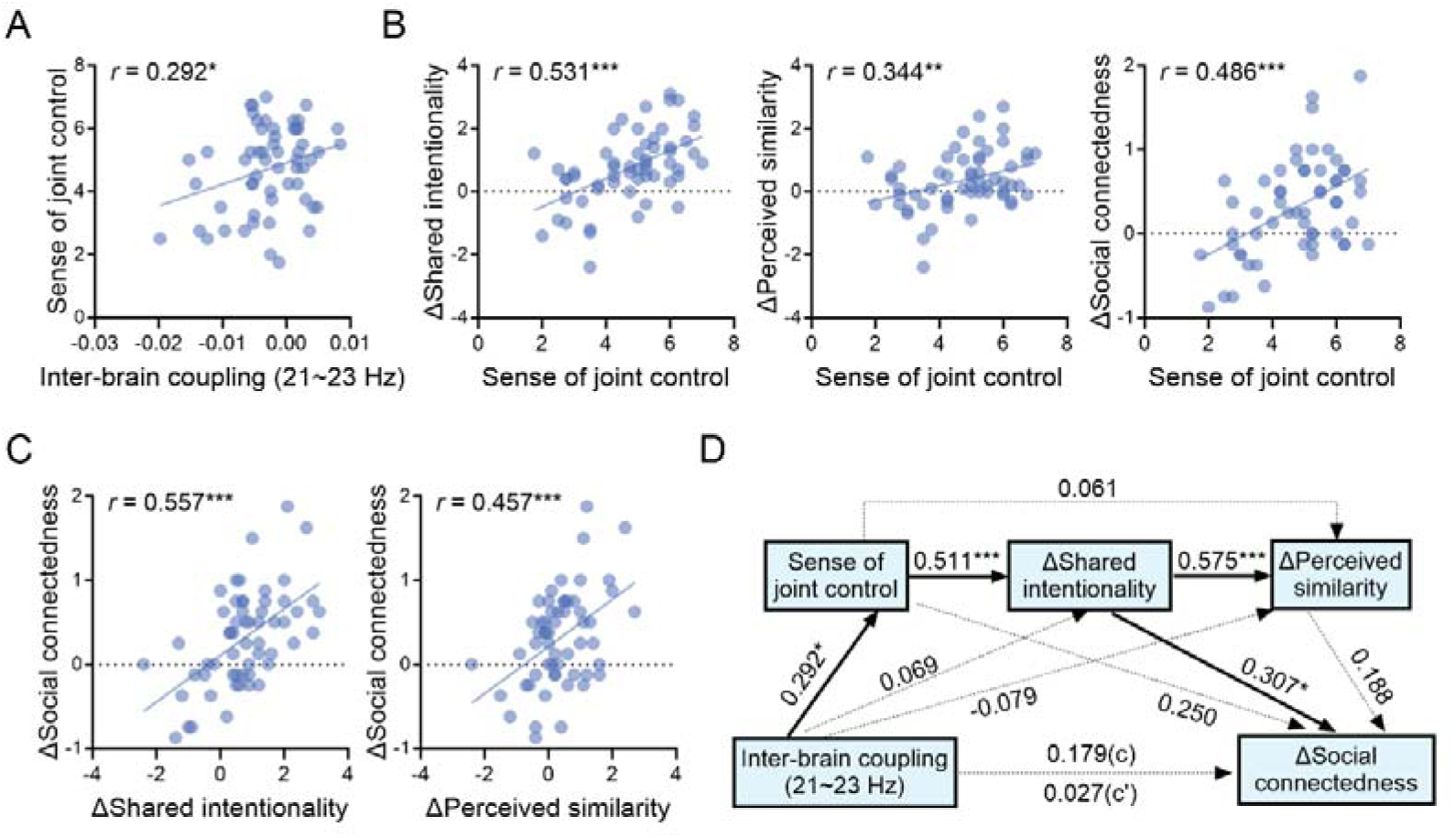
The relationship between inter-brain coupling at 21-23 Hz, sense of joint control, shared mental processes, and social connectedness. (A) Inter-brain coupling at 21-23 Hz was related to the sense of joint control. (B) Sense of joint control was associated with promoted shared intentionality (Δ Shared intentionality), promoted perceived similarity (Δ Perceived similarity), and social connectedness increase (Δ Social connectedness). (C) Promoted shared intentionality and perceived similarity predicted social connectedness increase. (D) The chain mediation result. Inter-brain coupling at 21-23 Hz affected the increase of social connectedness through sense of joint control and the improvement of shared intentionality. Path coefficients were standardized. Δ = post-training minus pre-training. * *p* < 0.05, ** *p* < 0.01, *** *p* < 0.001.

Based on these pairwise associations, we conducted an exploratory chain mediation analysis, with inter-brain coupling as the independent variable, sense of joint control, Δ Shared intentionality, and Δ Perceived similarity as potential mediators, and Δ Social connectedness as the dependent variable, respectively. This exploratory analysis revealed that inter-brain coupling at 21-23 Hz affected the increase of social connectedness through the sense of joint control and the improvement of shared intentionality, standardized indirect effect = 0.046 (bootstrap sample = 5000), *SE* = 0.031, 95% CI = [0.0003, 0.1212] (**Figure 5D**). Bootstrap indirect effects of other paths were not significant (**Table 1**).

**Table 1.** Standardized indirect effects of inter-brain coupling on Δ Social connectedness.

| Path | Effect | BootSE | 95%CI |
| --- | --- | --- | --- |
| Inter-brain coupling → Sense of joint control → $\Delta$ Social connectedness | 0.073 | 0.089 | [-0.036, 0.318] |
| Inter-brain coupling → $\Delta$ Shared intentionality → $\Delta$ Social connectedness | 0.021 | 0.038 | [-0.010, 0.185] |
| Inter-brain coupling → $\Delta$ Perceived similarity → $\Delta$ Social connectedness | -0.015 | 0.028 | [-0.080, 0.033] |
| **Inter-brain coupling → Sense of joint control → $\Delta$ Shared intentionality → $\Delta$ Social connectedness** | **0.046** | **0.031** | **[0.0003, 0.1212]** |
| Inter-brain coupling → Sense of joint control → $\Delta$ Perceived similarity → $\Delta$ Social connectedness | 0.003 | 0.009 | [-0.010, 0.028] |
| Inter-brain coupling → $\Delta$ Shared intentionality →<br>$\Delta$ Perceived similarity → $\Delta$ Social connectedness | 0.008 | 0.015 | [-0.020, 0.042] |
| Inter-brain coupling → Sense of control →<br>$\Delta$ Shared intentionality → $\Delta$ Perceived similarity →<br>$\Delta$ Social connectedness | 0.016 | 0.014 | [-0.007, 0.047] |

*Note.* SE = standard error, CI = confidence interval. $\Delta$ = post-training minus pre-training. Bootstrap sample = 5000.

## 4. Discussion

The current study used an EEG-based multi-person neurofeedback to investigate whether endogenously regulating inter-brain coupling can promote social connectedness. We found that dyads who received multi-person neurofeedback training (compared to a sham group) showed an increase in inter-brain coupling at 21-23 Hz. Task-related inter-brain coupling was correlated with dyads’ sense of social connectedness, while also improving the sense of joint control and shared mental processes (i.e., shared intentionality and perceived similarity), from pre-training to post-training. Finally, a chain mediation analysis revealed that inter-brain coupling at 21-23 Hz can predict increased social connectedness through the sense of joint control and the improvement of shared intentionality.

Our findings align with previous studies reporting a relationship between multi-person neurofeedback and social closeness and behavior ^38–40^. It is important to note that although the training target range was broadly set between 1-30Hz in the current study, offline analyses showed that the relevant frequency of interest was primarily concentrated at 21-23 Hz frequency band, within the beta frequency range. This may reflect a frequency-specific effect. This high-beta band (i.e., 21-23Hz) might be more responsive to the neurofeedback protocol used, as the brain’s intrinsic oscillatory properties may naturally favor this band under given training conditions. Previous studies have observed similar effects, suggesting that neurofeedback has frequency-specific predictions for social behaviors ^38^.

Importantly, we observed that real multi-person neurofeedback enhanced social connectedness and shared mental processes (shared intentionality and perceived similarity). Further, our chain mediation analysis suggests that inter-brain coupling regulated through multi-person neurofeedback may influence social connectedness through a sense of joint control and shared intentionality. These findings are consistent with prior studies linking changes in inter-brain coupling to cognitive and affective states ^18,19,58^, and cooperative performance and emotional resonance in dyads ^59–63^. This framework applied to our study: during multi-person neurofeedback, participants were asked to coordinate their thoughts with one another (i.e., pushing the avatar icons representing inter-brain coupling on the screen together with your mind). When individuals experience a sense of joint control, it fosters a shared understanding and coordination between them. This shared sense of agency may then translate into a shared intentionality to engage in mutual goals, which makes participants feel more aligned and connected with each other on cognitive and emotional levels ^25,64,65^. Consequently, this alignment of cognitive and emotional processes promoted their social connectedness, as individuals tend to offer social support more readily to those who resemble themselves ^66–69^. Overall, our findings highlight the intricate interplay between interpersonal dynamics, cognitive processes, and social outcomes in the context of neurofeedback-induced inter-brain coupling, underscoring the importance of shared experiences and intentions in promoting social connectedness.

In addition to its strengths, it is important to acknowledge several limitations in this study. First, the use of EEG-based neurofeedback imposes restrictions due to its limited spatial resolution. Future investigations could explore the integration of EEG with functional near-infrared spectroscopy (fNIRS), which offers better spatial resolution while keeping the naturalism. By combining these techniques, researchers can gain a more comprehensive understanding of both the temporal dynamics and spatial distribution of neural activity. Second, while multi-person neurofeedback showed promising group-level results, this study solely relied on self-reported measures of social connectedness. They may not fully capture the complexity and nuances of real-life social interactions and social connectedness. Therefore, future research should aim to complement self-reported measures with more objective assessments, such as behavioral observations or physiological markers of social connectedness.

## 5. Conclusion

In summary, our study not only confirms the feasibility of endogenous regulation of inter-brain coupling through multi-person neurofeedback but also adds to the existing body of literature by demonstrating that neurofeedback-regulated inter-brain coupling at 21-23 Hz can promote social connectedness, potentially through the sense of joint control and shared mental processes. Expanding on previous findings, our research underscores the important role that inter-brain coupling plays in shaping social interactions and relationships. Moreover, our study highlights the potential of regulating inter-brain coupling as a novel intervention strategy to optimize human social behaviors and foster healthier interpersonal connections.

## Author contributions

Conceptualization: XC, SD and YP. Methodology: PC and SD. Data collection and analysis: XC, RZ, ZS, FC. Writing—original draft: XC and YP. Writing—review and editing: XC, SD and YP.

## Acknowledgements

This work was supported by the Humanities and Social Sciences Research Project of the Ministry of Education of China (No. 24YJC190006) to X.C., the National Natural Science Foundation of China (Nos. 62207025 and 62337001), the Zhejiang Provincial Natural Science Foundation of China (No. LMS25C090002) to Y.P., and ERC-2023-CoG 101124369 to S.D.

## Data availability statement

The data that support the findings of this study are available on request from the corresponding author.

## References

1. Clark J.L., S.B. Algoe & M.C. Green. 2018. Social Network Sites and Well-Being: The Role of Social Connection. Curr. Dir. Psychol. Sci. 27: 32–37. 10.1177/0963721417730833

2. Lambert N.M., T.F. Stillman, J.A. Hicks, et al. 2013. To Belong Is to Matter: Sense of Belonging Enhances Meaning in Life. Personal. Soc. Psychol. Bull. 39: 1418–1427. 10.1177/0146167213499186

3. Christiansen J., P. Qualter, K. Friis, et al. 2021. Associations of loneliness and social isolation with physical and mental health among adolescents and young adults. Perspect. Public Health 141: 226–236. 10.1177/17579139211016077

4. Freak-Poli R., J. Ryan, J.T. Neumann, et al. 2021. Social isolation, social support and loneliness as predictors of cardiovascular disease incidence and mortality. BMC Geriatr. 21: 711. 10.1186/s12877-021-02602-2

5. Humphrey A., E. March, A.P. Lavender, et al. 2022. Buffering the Fear of COVID-19: Social Connectedness Mediates the Relationship between Fear of COVID-19 and Psychological Wellbeing. Behav. Sci. (Basel). 12: 86. 10.3390/bs12030086

6. Kim M.J. & S. Sul. 2023. On the relationship between the social brain, social connectedness, and wellbeing. Front. Psychiatry 14: 1112438. 10.3389/fpsyt.2023.1112438

7. Cheng X., S. Wang, B. Guo, et al. 2024. How self-disclosure of negative experiences shapes prosociality? Soc. Cogn. Affect. Neurosci. 19: nsae003. 10.1093/scan/nsae003

8. Xie E., Q. Yin, K. Li, et al. 2021. Sharing Happy Stories Increases Interpersonal Closeness: Interpersonal Brain Synchronization as a Neural Indicator. eneuro 8: ENEURO.0245-21.2021. 10.1523/ENEURO.0245-21.2021

9. Jiang J., B. Dai, D. Peng, et al. 2012. Neural synchronization during face-to-face communication. J. Neurosci. 32: 16064–16069. 10.1523/JNEUROSCI.2926-12.2012

10. Jiang J., C. Chen, B. Dai, et al. 2015. Leader emergence through interpersonal neural synchronization. Proc. Natl. Acad. Sci. U. S. A. 112: 4274–4279. 10.1073/pnas.1422930112

11. Pan Y., S. Dikker, P. Goldstein, et al. 2020. Instructor-learner brain coupling discriminates between instructional approaches and predicts learning. Neuroimage 211: 116657. 10.1016/j.neuroimage.2020.116657

12. Zhu Y., V. Leong, Y. Hou, et al. 2022. Instructor–learner neural synchronization during elaborated feedback predicts learning transfer. J. Educ. Psychol. 114: 1427–1441. 10.1037/edu0000707

13. Hu Y., Y. Pan, X. Shi, et al. 2018. Inter-brain synchrony and cooperation context in interactive decision making. Biol. Psychol. 133:. 10.1016/j.biopsycho.2017.12.005

14. Tang H., X. Mai, S. Wang, et al. 2015. Interpersonal brain synchronization in the right temporo-parietal junction during face-to-face economic exchange. Soc. Cogn. Affect. Neurosci. 10.1093/scan/nsv092

15. Cheng X., Y. Pan, Y. Hu, et al. 2019. Coordination Elicits Synchronous Brain Activity Between Co-actors: Frequency Ratio Matters. Front. Neurosci. 13: 1071. 10.3389/fnins.2019.01071

16. Long Y., M. Zhong, R. Aili, et al. 2023. Transcranial direct current stimulation of the right anterior temporal lobe changes interpersonal neural synchronization and shared mental processes. Brain Stimul. 16: 28–39. 10.1016/j.brs.2022.12.009

17. Novembre G., G. Knoblich, L. Dunne, et al. 2017. Interpersonal synchrony enhanced through 20 Hz phase-coupled dual brain stimulation. Soc. Cogn. Affect. Neurosci. 12: 662–670. 10.1093/scan/nsw172

18. Liu L., Y. Zhang, Q. Zhou, et al. 2020. Auditory–Articulatory Neural Alignment between Listener and Speaker during Verbal Communication. Cereb. Cortex 30: 942–951. 10.1093/cercor/bhz138

19. Zheng L., W. Liu, Y. Long, et al. 2020. Affiliative bonding between teachers and students through interpersonal synchronisation in brain activity. Soc. Cogn. Affect. Neurosci. 15: 97–109. 10.1093/scan/nsaa016

20. Iacoboni M. 2005. Neural mechanisms of imitation. Curr. Opin. Neurobiol. 15: 632–637. 10.1016/j.conb.2005.10.010

21. Keysers C. & V. Gazzola. 2014. Hebbian learning and predictive mirror neurons for actions, sensations and emotions. Philos. Trans. R. Soc. B Biol. Sci. 369: 20130175. 10.1098/rstb.2013.0175

22. Rizzolatti G. & C. Sinigaglia. 2016. The mirror mechanism: a basic principle of brain function. Nat. Rev. Neurosci. 17: 757–765. 10.1038/nrn.2016.135

23. Kirschner S. & M. Tomasello. 2010. Joint music making promotes prosocial behavior in 4-year-old childrenLLL. Evol. Hum. Behav. 31: 354–364. 10.1016/j.evolhumbehav.2010.04.004

24. Mazzurega M., F. Pavani, M.P. Paladino, et al. 2011. Self-other bodily merging in the context of synchronous but arbitrary-related multisensory inputs. Exp. Brain Res. 213: 213–221. 10.1007/s00221-011-2744-6

25. Reddish P., R. Fischer & J. Bulbulia. 2013. Let’s Dance Together: Synchrony, Shared Intentionality and Cooperation. PLoS One 8: e71182. 10.1371/journal.pone.0071182

26. Hu Y., Y. Hu, X. Li, et al. 2017. Brain-to-brain synchronization across two persons predicts mutual prosociality. Soc. Cogn. Affect. Neurosci. 12:. 10.1093/scan/nsx118

27. Valdesolo P., J. Ouyang & D. DeSteno. 2010. The rhythm of joint action: Synchrony promotes cooperative ability. J. Exp. Soc. Psychol. 46: 693–695. 10.1016/j.jesp.2010.03.004

28. Alegria A.A., M. Wulff, H. Brinson, et al. 2017. RealLtime fMRI neurofeedback in adolescents with attention deficit hyperactivity disorder. Hum. Brain Mapp. 38: 3190–3209. 10.1002/hbm.23584

29. Jurewicz K., K. Paluch, E. Kublik, et al. 2018. EEG-neurofeedback training of beta band (12–22 Hz) affects alpha and beta frequencies – A controlled study of a healthy population. Neuropsychologia 108: 13–24. 10.1016/j.neuropsychologia.2017.11.021

30. Agnoli S., M. Zanon, S. Mastria, et al. 2018. Enhancing creative cognition with a rapid right-parietal neurofeedback procedure. Neuropsychologia 118: 99–106. 10.1016/j.neuropsychologia.2018.02.015

31. Coben R., M. Middlebrooks, H. Lightstone, et al. 2018. Four Channel Multivariate Coherence Training: Development and Evidence in Support of a New Form of Neurofeedback. Front. Neurosci. 12: 729. 10.3389/fnins.2018.00729

32. Loriette C., C. Ziane & S. Ben Hamed. 2021. Neurofeedback for cognitive enhancement and intervention and brain plasticity. Rev. Neurol. (Paris). 177: 1133–1144. 10.1016/j.neurol.2021.08.004

33. Gruzelier J.H. 2014. EEG-neurofeedback for optimising performance. I: A review of cognitive and affective outcome in healthy participants. Neurosci. Biobehav. Rev. 44: 124–141. 10.1016/j.neubiorev.2013.09.015

34. Gruzelier J.H. 2014. EEG-neurofeedback for optimising performance. III: A review of methodological and theoretical considerations. Neurosci. Biobehav. Rev. 44: 159–182. 10.1016/j.neubiorev.2014.03.015

35. Eschmann K.C.J., R. Bader & A. Mecklinger. 2020. Improving episodic memory: Frontal-midline theta neurofeedback training increases source memory performance. Neuroimage 222: 117219. 10.1016/j.neuroimage.2020.117219

36. Conti D., L. Celebre, N. Girone, et al. 2021. An intensive neurofeedback alpha-training to improve sleep quality and stress modulation in health-care workers during the COVID-19 pandemic: A pilot study. Eur. Psychiatry 64: S263–S263. 10.1192/j.eurpsy.2021.705

37. Chen P., S. Hendrikse, K. Sargent, et al. 2021. Hybrid Harmony: A multi-person neurofeedback application for interpersonal synchrony. Front. Neuroergonomics 2: 687108. 10.3389/fnrgo.2021.687108

38. Dikker S., G. Michalareas, M. Oostrik, et al. 2021. Crowdsourcing neuroscience: Inter-brain coupling during face-to-face interactions outside the laboratory. Neuroimage 227: 117436. 10.1016/j.neuroimage.2020.117436

39. Duan L., W.J. Liu, R.N. Dai, et al. 2013. Cross-brain neurofeedback: Scientific concept and experimental platform. PLoS One 8: e64590. 10.1371/journal.pone.0064590

40. Müller V., D. Perdikis, M.A. Mende, et al. 2021. Interacting brains coming in sync through their minds: an interbrain neurofeedback study. Ann. N. Y. Acad. Sci. 1500: 48–68. 10.1111/nyas.14605

41. Cheng X., X. Li & Y. Hu. 2015. Synchronous brain activity during cooperative exchange depends on gender of partner: A fNIRS-based hyperscanning study. Hum. Brain Mapp. 36: 2039–2048. 10.1002/hbm.22754

42. Di Bartolomeo G. & S. Papa. 2019. The Effects of Physical Activity on Social Interactions: The Case of Trust and Trustworthiness. J. Sports Econom. 20: 50–71. 10.1177/1527002517717299

43. Epley N., M. Kardas, X. Zhao, et al. 2022. Undersociality: miscalibrated social cognition can inhibit social connection. Trends Cogn. Sci. 26: 406–418. 10.1016/j.tics.2022.02.007

44. Bastian B., J. Jetten & L.J. Ferris. 2014. Pain as Social Glue: Shared Pain Increases Cooperation. Psychol. Sci. 25: 2079–2085. 10.1177/0956797614545886

45. Dumas G., J. Nadel, R. Soussignan, et al. 2010. Inter-brain synchronization during social interaction. PLoS One 5: e12166. 10.1371/journal.pone.0012166

46. Puccinelli N.M. & L. Tickle-Degnen. 2004. Knowing too much about others: Moderators of the relationship between eavesdropping and rapport in social interaction. J. Nonverbal Behav. 28: 223–243. 10.1007/s10919-004-4157-8

47. Batson C.D. 2010. Empathy-induced altruistic motivation. In Prosocial motives, emotions, and behavior: The better angels of our nature. 15–34. Washington: American Psychological Association. 10.1037/12061-001

48. Dikker S., L. Wan, I. Davidesco, et al. 2017. Brain-to-brain synchrony tracks real-world dynamic group interactions in the classroom. Curr. Biol. 27: 1375– 1380. 10.1016/j.cub.2017.04.002

49. Ayrolles A., F. Brun, P. Chen, et al. 2021. HyPyP: a Hyperscanning Python Pipeline for inter-brain connectivity analysis. Soc. Cogn. Affect. Neurosci. 16: 72–83. 10.1093/scan/nsaa141

50. Berens P. 2009. CircStatL: A MATLAB toolbox for circular statistics. J. Stat. Softw. 31: 1–21. 10.18637/jss.v031.i10

51. Pérez A., M. Carreiras & J.A. Duñabeitia. 2017. Brain-To-brain entrainment: EEG interbrain synchronization while speaking and listening. Sci. Rep. 10.1038/s41598-017-04464-4

52. Wikström V., K. Saarikivi, M. Falcon, et al. 2022. Inter-brain synchronization occurs without physical co-presence during cooperative online gaming. Neuropsychologia 174: 108316. 10.1016/j.neuropsychologia.2022.108316

53. Delorme A. & S. Makeig. 2004. EEGLAB: an open source toolbox for analysis of single-trial EEG dynamics including independent component analysis. J. Neurosci. Methods 134: 9–21. 10.1016/j.jneumeth.2003.10.009

54. Mognon A., J. Jovicich, L. Bruzzone, et al. 2011. ADJUST: An automatic EEG artifact detector based on the joint use of spatial and temporal features. Psychophysiology 48: 229–240. 10.1111/j.1469-8986.2010.01061.x

55. Jahangir M., Syed Tanweer, Rizwan-ur-Rehman, et al. 2017. Spectral and Spatial Feature Extraction of Electroencephalographic (EEG) Data Using Independent Component Analysis (ICA). J. Basic Appl. Sci. 13: 104–113. 10.6000/1927-5129.2017.13.18

56. Mu Y., C. Guo & S. Han. 2016. Oxytocin enhances inter-brain synchrony during social coordination in male adults. Soc. Cogn. Affect. Neurosci. 11: 1882–1893. 10.1093/scan/nsw106

57. Pérez A., G. Dumas, M. Karadag, et al. 2019. Differential brain-to-brain entrainment while speaking and listening in native and foreign languages. Cortex 111: 303–315. 10.1016/j.cortex.2018.11.026

58. Dai B., C. Chen, Y. Long, et al. 2018. Neural mechanisms for selectively tuning in to the target speaker in a naturalistic noisy situation. Nat. Commun. 9: 1–12. 10.1038/s41467-018-04819-z

59. Hu Y., X. Cheng, Y. Pan, et al. 2022. The intrapersonal and interpersonal consequences of interpersonal synchrony. Acta Psychol. (Amst). 224: 103513. 10.1016/j.actpsy.2022.103513

60. Knoblich G., S. Butterfill & N. Sebanz. 2011. Psychological Research on Joint Action. In 59–101. 10.1016/B978-0-12-385527-5.00003-6

61. Pan Y., X. Cheng, Z. Zhang, et al. 2017. Cooperation in lovers: An fNIRS-based hyperscanning study. Hum. Brain Mapp. 38: 831–841. 10.1002/hbm.23421

62. Reinero D.A., S. Dikker & J.J. Van Bavel. 2021. Inter-brain synchrony in teams predicts collective performance. Soc. Cogn. Affect. Neurosci. 16: 43–57. 10.1093/scan/nsaa135

63. Schippers M.B., A. Roebroeck, R. Renken, et al. 2010. Mapping the information flow from one brain to another during gestural communication. Proc. Natl. Acad. Sci. 107: 9388–9393. 10.1073/pnas.1001791107

64. Barraza P., A. Pérez & E. Rodríguez. 2020. Brain-to-Brain Coupling in the Gamma-Band as a Marker of Shared Intentionality. Front. Hum. Neurosci. 14: 295. 10.3389/fnhum.2020.00295

65. Fishburn F.A., V.P. Murty, C.O. Hlutkowsky, et al. 2018. Putting our heads together: interpersonal neural synchronization as a biological mechanism for shared intentionality. Soc. Cogn. Affect. Neurosci. 13: 841–849. 10.1093/scan/nsy060

66. Kehayes I.-L.L., S.P. Mackinnon, S.B. Sherry, et al. 2017. Similarity in Romantic Couples’ Drinking Motivations and Drinking Behaviors. Subst. Abus. 38: 488–492. 10.1080/08897077.2017.1355869

67. Martin B., A. Giersch, C. Huron, et al. 2013. Temporal event structure and timing in schizophrenia: Preserved binding in a longer “now.” Neuropsychologia 51: 358–371. 10.1016/j.neuropsychologia.2012.07.002

68. Westmaas J.L. & R.C. Silver. 2006. The Role of Perceived Similarity in Supportive Responses to Victims of Negative Life Events. Personal. Soc. Psychol. Bull. 32: 1537–1546. 10.1177/0146167206291874

69. Whitmore C.B. & J.C. Dunsmore. 2014. Trust Development: Testing a New Model in Undergraduate Roommate Relationships. J. Genet. Psychol. 175: 233–251. 10.1080/00221325.2013.869533

